# Evidence for Increasingly Variable Drought Conditions in the United States Since 1895

**DOI:** 10.1101/030031

**Authors:** Sierra Rayne, Kaya Forest

## Abstract

Annual and summertime trends towards increasingly variable values of the Palmer Drought Severity Index (PDSI) over a sub-decadal period (five years) were investigated within the contiguous United States between 1895 and the present. For the contiguous U.S. as a whole, there is a significant increasing trend in the five-year running minimum-maximum ranges for the annual PDSI (_a_PDSI_5yr(min| max range))_. During this time frame, the average _a_PDSI_5yr(min|max range)_ has increased by about one full unit, indicating a substantial increase is drought variability over short time scales across the United States. The end members of the running _a_PDSI_5yr(min|max range)_ highlight even more rapid changes in the drought index variability within the past 120 years. This increasing variability in the _a_PDSI_5yr(min|max range)_ is driven primarily by changes taking place in the Pacific and Atlantic Ocean coastal climate regions, climate regions which collectively comprise one-third the area of the contiguous U.S. Overall, interannual drought patterns are becoming more extreme and difficult to predict, posing a challenge to agricultural and other water-resource related planning efforts.

## 1 Introduction

The severe ecological and socio-economic impacts of droughts have led many to consider the potential effects of climate change on the frequency, duration, and intensity of these phenomena (Kallis 2008). At a global scale, drought is expected to increase in frequency and intensity during the twenty-first century (Dai et al. 2004; Briffa et al. 2009; Burke et al. 2006; Sheffield and Wood 2008a; Dai 2011a), although there is modest disagreement as to the magnitude and direction of any recent trends (Dai 2011b; Sheffield and Wood 2008b). Furthermore, in addition to established anthropogenic forcings, various natural climate cycles (e.g., El Nino-Southern Oscillation [ENSO], Atlantic Multi-Decadal Oscillation [AMO], Pacific Decadal Oscillation [PDO]) can also complicate time trend analyses of drought conditions (Cole et al. 2002; Cook et al. 2010; Dai 2011a; Hidalgo 2004; McCabe et al. 2004; Seager and Vecchi 2010; Woodhouse and Overpeck 1998).

In response to these challenges, various measures for assessing drought conditions have been developed (Dai 2011b; Heim 2002; Kallis 2008; Keyantash and Dracup 2002; Piechota and Dracup 1996), but the Palmer suite of indices (e.g., Palmer Drought Severity Index [PDSI]) remains in wide application. Whereas flaws in the PDSI and related approaches for drought determination and prediction have been broadly discussed (Alley 1984; Andreadis et al. 2005; Briffa et al. 2009; Dai 2011b; Dai et al. 2004; Heim 2002; Hoerling et al. 2013; Kallis 2008; Keyantash and Dracup 2002; Piechota and Dracup 1996; Sheffield et al. 2012), the general consensus appears to be that PDSI-related metrics still offer primary utility for climate change studies.

In the United States, prior work has shown historical trends and predictions toward increasingly frequent and severe droughts and a generally drier hydroclimate in the western and southwestern regions (Andreadis and Lettenmaier 2006; Cayan et al. 2010; Cook et al. 2004; Seager et al. 2007, 2013; Schwalm et al. 2012; Sheffield and Wood 2008b) and no clear changes, or reduced drought conditions, in the rest of the nation (Andreadis and Lettenmaier 2006; Dai et al. 2004; Seager et al. 2009). Recent modeling work suggests the the Great Plains region will be in semi-permanent severe drought over the coming century (Hoerling et al. 2013).

Overall, current observable trends for regional drought indices in the United States are in agreement with climate models that associate substantial portions of the changes to anthropogenic causes, thereby warranting additional mechanistic and empirical studies. In the current work, we extend these investigations to the question of whether drought conditions in the United States are becoming more variable over sub-decadal periods of time.

## 2. Datasets and Methods

Data was obtained from the National Centers for Environmental Information of the National Oceanic and Atmospheric Administration (https://www.ncdc.noaa.gov/cag/). Individual climate regions (Karl and Koss 1984) under more detailed examination comprise the following states: Northwest (Idaho, Oregon, and Washington); West (California and Nevada); Northeast (Connecticut, Delaware, Maine, Maryland, Massachusetts, New Hampshire, New Jersey, New York, Pennsylvania, Vermont, and Rhode Island).

Statistical analyses of climate data were conducted using the nonparametric Mann-Kendall test for the trend and the nonparametric Sen’s method for the magnitude of the trend (Kendall 1975; Mann 1945; Salmi et al. 2002) within the R Project for Statistical Computing environment (R Core Team 2012). Parametric linear regressions and non-parametric Spearman rank correlations were performed with KyPlot (v.2.0b15) (Yoshioka 2002).

## 3. Results and Discussion

Temporal alterations in drought variability can be investigated by considering trends in the five-year running minimum-maximum ranges for the annual Palmer Drought Severity Index (_a_PDSI_5yr(min|max range))_ over this time frame. Such metrics capture cases whereby widely varying drought conditions (e.g., an extreme dry year is followed by an extreme wet year) are becoming more prevalent. For the contiguous U.S., there is a significant (p=0.076/0.14/0.18; parametric linear/Mann-Kendall/Spearman rank; m=0.0085±0.0047 [SE] PDSI units/yr) increasing trend in the _a_PDSI_5yr(min|max range)_ between 1895 and 2014 (Figure 1). No corresponding significant trend is present for the _a_PDSI_5yr(min|max range)_ residuals, supporting the assignment of a linear trend analysis. During this period, the average _a_PDSI_5yr(min|max range)_ has increased by about one full unit, representing a substantial increase is drought variability over short time scales across the United States. Consequently, during a half-decade, interannual drought patterns are becoming more extreme and difficult to predict, posing a challenge to agricultural and other water-resource related planning efforts.

**Figure 1.**
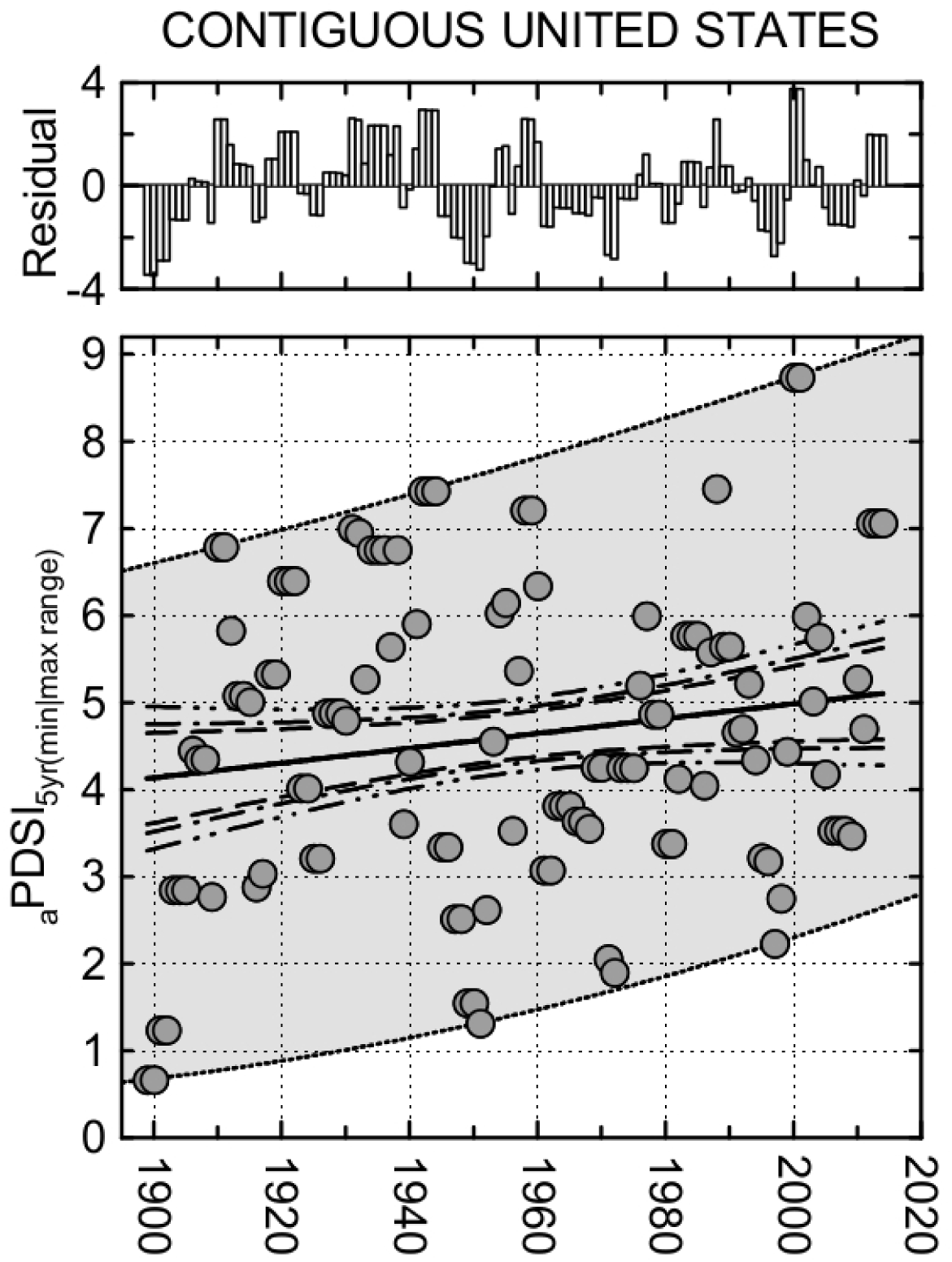
Five-year running minimum-maximum ranges for the annual Palmer Drought Severity Index (_a_PDSI_5yr(min|max range))_ over the contiguous United States between 1895 and 2014. Dashed, dashed-dot, and dashed-dot-dot lines are upper and lower 90%, 95%, and 99% confidence intervals about the regression line (solid). Inset shows residuals for the linear regression in the main figure. Dotted lines and shading encompass the minimum-maximum envelope in _a_PDSI_5yr(min|max range)_ as it evolves over time.

**Figure 2.**
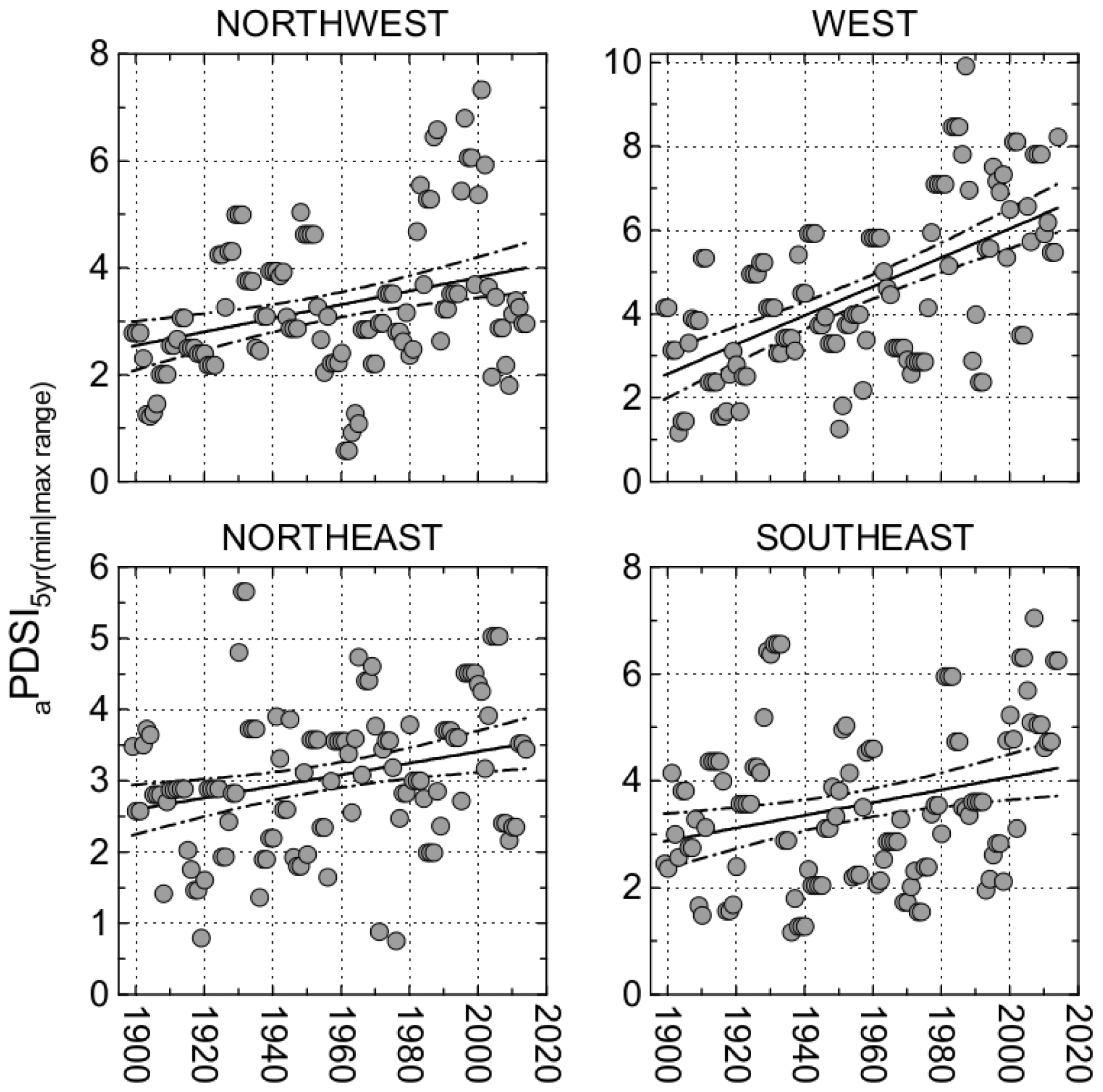
Five-year running minimum-maximum ranges for the annual Palmer Drought Severity Index (_a_PDSI_5yr(min|max range))_ in the Northwest, West, Northeast, and Southeast climate regions between 1895 and 2014. The dashed-dot lines are upper and lower 95% confidence intervals about the regression line (solid).

The end members of the running _a_PDSI_5yr(min|max range)_ highlight even more rapid changes in the drought index variability. In the late 1890s and early 1900s, _a_PDSI_5yr(min|max range)_ on the order of 1.0 -- indicating very low inter-annual variability (i.e., climatic stability) -- were evident. By mid-century, the lowest _a_PDSI_5yr(min|max range)_ had increased up to 1.5 PDSI units, reaching two PDSI units by the early 1970s. Since 1999, the lowest _a_PDSI5_yr(min|max range)_ was 3.5 between 2006 and 2009. At the high end of the scale, _a_PDSI_5yr(min|max range)_ of less than 7 PDSI units were the maximum occurrences in the early 20th century. By mid-century, the variability had increased to ~7.5 PDSI units, and in 2000/2001 the index reached a record-high 8.74 PDSI units when the U.S. as a whole abruptly swung from an extreme wet year in 1997 (PDSI=4.30) to an extreme drought year (PDSI=-4.44) just three years later in 2000. This 5-year drought variability from 1997 through 2000 far exceeded any prior maximum variation in the historical record, but also fits into the upper envelope trendline for maximum _a_PDSI5_yr(min|max range)_ since 1895.

The increasing variability in the _a_PDSI_5yr(min|max range)_ is driven primarily by changes taking place in the Pacific and Atlantic Ocean coastal climate regions (Northwest, West, Northeast, and Southeast). Highly significant increasing trends in the _a_PDSI5_yr(min|max range)_ are evident since 1895 in the Northwest (p=0.0004/0.0006/0.0006; m=0.013±0.003 PDSI units/yr), West (p=2×10^−12^/2×10^−16^/2×10^−9^; m=0.035±0.004 PDSI units/yr), Northeast (p=0.003/0.002/0.003; m=0.0081±0.0027 PDSI units/yr), and Southeast (p=0.003/0.003/0.006; m=0.011±0.004 PDSI units/yr) regions. Analogous significant trends towards increasingly variable five-year PDSI ranges are present for these climate regions during the summer months between 1895 and 2015 (_s_PDSI5_yr(min|max range)_; Figure 3): Northwest, p=0.0006/0.0006/0.0008,m=0.014±0.004 PDSI units/yr; West, p=6×10^−11^/2×10^−16^/4×10^−9^,m=0.039±0.005 PDSI units/yr; Northeast, p=4×10^−8^/<10^−16^/4×10^−8^,m=0.018±0.003 PDSI units/yr; and Southeast, p=0.003/0.047/0.02,m=0.015±0.005 PDSI units/yr.

**Figure 3.**
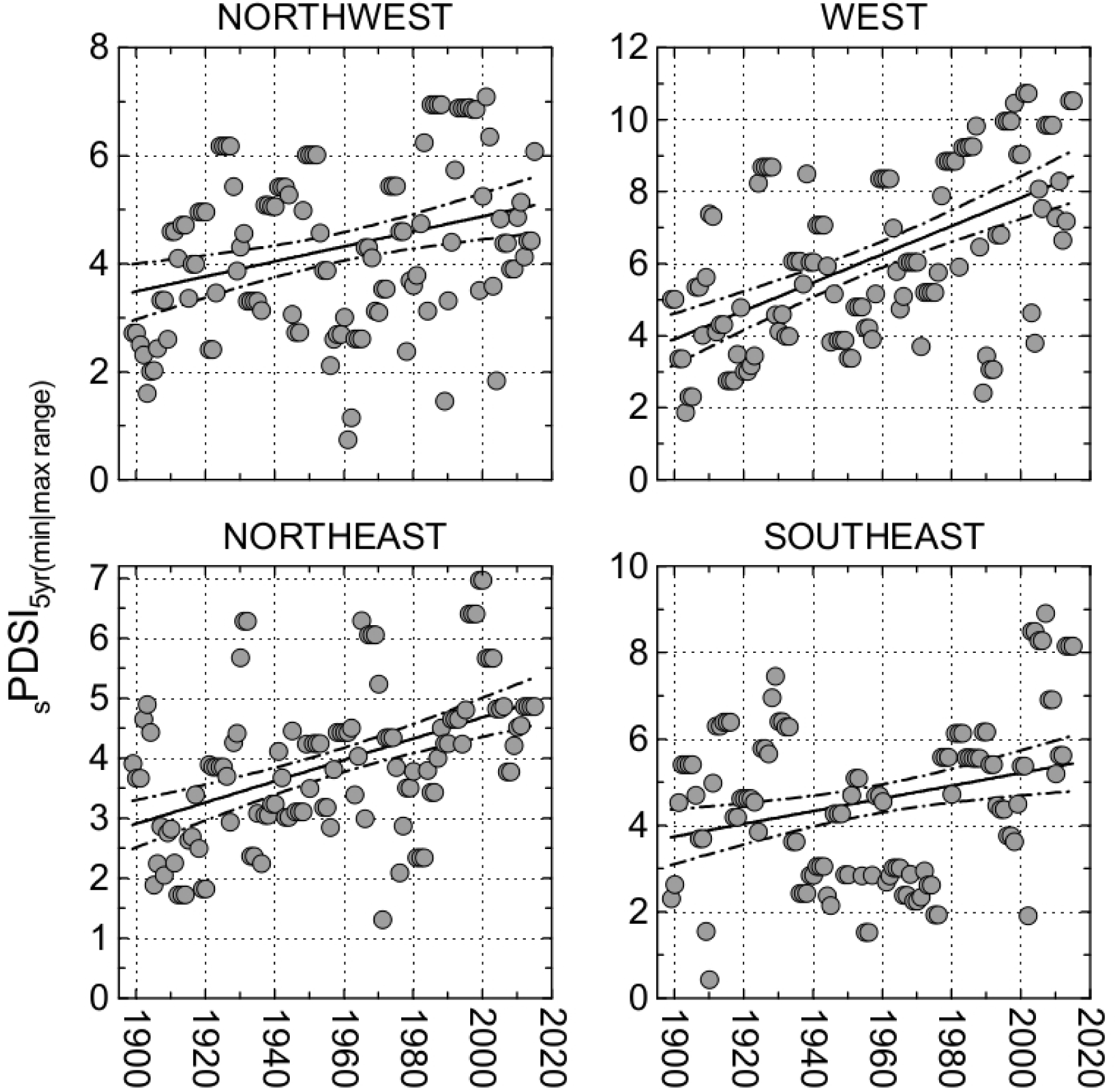
Five-year running minimum-maximum ranges for the summertime (June to August) Palmer Drought Severity Index (_s_PDSI_5yr(min|max range)_) in the Northwest, West, Northeast, and Southeast climate regions between 1895 and 2014. The dashed-dot lines are upper and lower 95% confidence intervals about the regression line (solid).

Together, these climate regions comprise one-third the area of the contiguous U.S. The patterns over time display strong correlations with respective regional temperature trends, as well as with Northern Hemisphere ocean temperature increases since the late 19th century, which collectively have been linked to anthropogenic drivers (IPCC 2013, 2014a, 2014b). The effects of the cool (1890 to 1924 and 1947 to 1976) and warm (1925 to 1946 and 1977 to the mid-1990s) Pacific Decadal Oscillation (PDO) regimes (Trenberth 1990; Latif and Barnett 1994; Trenberth and Hurrell 1994; Minobe 1997; Zhang et al. 1997; Barnett et al. 1999; Nigam et al. 1999) show up clearly in the Northwest climate region _a_PDSI_5yr(min|max range)_ and _s_PDSI_5yr(min|max range)_ time trends, with the largest _a/s_PDSI_5yr(min|max range)_ taking place during warm PDO phases. However, only modest, positive correlations we obtain between the 1900–2014 annual average and summertime PDO indices (http://research.jisao.washington.edu/pdo/PDO.latest) and the corresponding _a_PDSI5_yr(min|max range)_ (r=+0.34; p=0.0002) and _s_PDSI5_yr(min|max range)_ (r=+0.12; p=0.20) suggests this natural climatic variability plays only a minor role in the observed increasing drought variability for this area.

In conclusion, drought variability within five-year running period sub-decadal time scales is rapidly increasing in the coastal climate regions of the United States and for the contiguous nation as a whole. Further mechanistic studies are warranted to determine the potential causal underpinnings of these trends, but at present, known natural cycles do not appear capable of explaining the changes.

